# Synergistic cefiderocol-containing antibiotic combinations active against highly drug-resistant *Acinetobacter baumannii* patient isolates with diverse resistance mechanisms

**DOI:** 10.1101/2025.04.22.649919

**Authors:** Justin Halim, Andrew P. Keane, Jeannete Bouzo, Tope Aderibigbe, Jessica A. Chicola, Katie T. Nolan, Keertana Jonnalgadda, Jason X. Tran, Valerie J. Carabetta

## Abstract

*Acinetobacter baumannii* is a nosocomial pathogen notorious for rapidly acquiring resistance to nearly all antibiotics, including last-line agents. Cefiderocol (FDC) is a novel siderophore cephalosporin antibiotic with *in vitro* activity against *A. baumannii* that is now used clinically, but resistance is emerging. There is limited data regarding FDC-containing antibiotic combinations with synergistic activity against *A. baumannii*, which may increase the potency of the drug and reduce resistance development. In this study, we evaluated the activity of FDC alone and in combination with other antibiotics against 21 extensively drug-resistant (XDR) and pandrug-resistant (PDR) *A. baumannii* clinical isolates. FDC possesses strong *in vitro* activity and was highly effective in combination with ceftazidime-avibactam (CAZ-AVI) and sulbactam-durlobactam (SUL-DUR), as well as amikacin, doxycycline, and sulbactam. We also identified several FDC-containing combinations effective against a metallo-β-lactamase (MBL)-producing strain. Previously, we performed whole genome sequencing on selected strains in our collection, allowing for the identification of resistance genes and mechanisms associated with susceptibility patterns and synergy outcomes. These findings highlight promising FDC-based combinations for treating highly drug-resistant *A. baumannii* infections and provide insight into the genetic basis of FDC resistance and response.

**Author Summary:** *Acinetobacter baumannii* is a frequent cause of hospital-acquired pneumonia, and bloodstream and wound infections in vulnerable populations, such as those who are critically ill or immunocompromised. Infections caused by *A. baumannii* are a growing problem in hospitals around the world, especially because of increasing resistance to almost all available antibiotics. Physicians have few treatment options available, which leads to high mortality rates. One newer antibiotic, cefiderocol, has shown promise against these infections, but resistance to this drug is already emerging. In our study, we looked for combinations of cefiderocol with other traditional and newly available antibiotics that work better together than alone. We tested these combinations against over 20 highly drug-resistant strains isolated from patients, including one strain that produces a strong resistance enzyme called NDM-1. We found that pairing cefiderocol with certain drugs, especially ceftazidime-avibactam, sulbactam-durlobactam, amikacin, and doxycycline, often worked significantly better than using cefiderocol alone. These combinations were able to prevent the growth of even the most resistant strains in our collection. Our work highlights new potential treatment options that physicians may use in the future to help patients with these dangerous, life-threatening infections, especially when standard antibiotics are no longer effective.

## 1. Introduction

*Acinetobacter baumannii* is an aerobic, Gram-negative coccobacillus that, while typically of low virulence, is a common cause of hospital-acquired infections, particularly in immunocompromised patients and those undergoing intensive care, mechanical ventilation, or catheterization [1,2]. *A. baumannii* infections demonstrate high mortality rates, ranging from 40% to 80% [3,4]. Resistance develops rapidly via mechanisms such as expression of efflux pumps, aminoglycoside modifying enzymes (AMEs) and β-lactamases, and mutations in drug targets, such as penicillin-binding proteins (PBPs) [5]. The emergence of multidrug-resistant (MDR), extensively drug-resistant (XDR), and pandrug-resistant (PDR) strains is now a global threat [5,6]. In particular, carbapenem-resistant *A. baumannii* (CRAB), defined as resistance to imipenem and meropenem (MIC ≥8 μg/mL), is often resistant to last-line antibiotics including carbapenems, colistin, and tigecycline [7–9]. As of 2019, the Center for Disease Control and Prevention categorizes CRAB infections as an urgent public health threat [10].

Cefiderocol (FDC), a novel siderophore cephalosporin, was recently approved by the U.S. Food and Drug Administration for hospital- and ventilator-associated pneumonia (VAP) caused by Gram-negative species, including MDR *Pseudomonas aeruginosa, A. baumannii,* and *Enterobacteriaceae* [11]. Structurally related to ceftazidime and cefepime, FDC also features a catechol moiety that enables iron-mediated uptake into bacterial cells, bypassing entry through porins [12]. This novel mechanism increases uptake despite porin mutations or efflux mechanisms, and the large structure of FDC confers stability against β-lactamases of all Ambler classes, including AmpC and the extended spectrum β-lactamases (ESBLs) [12]. FDC demonstrates both *in vitro* activity against CRAB isolates and clinical efficacy in patients with *A. baumannii* infections [13–15]. However, clinical trial data have been mixed. The CREDIBLE-CR trial found similar infection resolution but higher mortality with FDC compared to best available therapy [16]. The APEKS-NP trial, in contrast, showed FDC to be non-inferior to meropenem for nosocomial pneumonia [17]. Due to these mixed results, the Infectious Diseases Society of America (IDSA) recommends using FDC for CRAB only when alternatives are unavailable, and always in combination [18]. FDC resistance is increasingly reported, especially in strains expressing OXA-type or metallo-β-lactamases (MBLs), such as the New Delhi MBL, NDM-1 [19]. MBLs are of particular concern, as they confer resistance to nearly all β-lactam drugs and are associated with highly resistant phenotypes [19].

Antimicrobial synergism, defined as when two drugs yield greater activity together than individually, can improve efficacy and reduce resistance development [20]. Data on FDC-based combinations against *A. baumannii* remain limited. Here, we evaluated the *in vitro* activity of FDC in combination with other antibiotics against 21 XDR and PDR *A. baumannii* clinical isolates. We evaluated FDC in combination with 17 standard-of-care antibiotics, as well as sulbactam-durlobactam (SUL-DUR), eravacycline, and omadacycline, which were recently introduced to the market with noted *in vitro* activity against *A. baumannii*. Prior whole genome sequencing (WGS) found that our collection contained diverse beta-lactamases but lacked MBLs.^21^ To assess synergy against MBL-harboring strains, we also included a known *NDM-1*-producing strain. Our aim was to identify synergistic FDC-based combinations and assess the contributions of beta-lactamase production as potential FDC resistance mechanisms. We identified multiple promising FDC combinations that exhibited synergy against most strains tested.

## 2. Results

### 2.1. Determination of susceptibilities of A. baumannii clinical isolates to FDC

We evaluated the susceptibility of 21 previously characterized XDR and PDR *A. baumannii* isolates (strains M1–M22) to FDC via broth microdilution [21]. Fourteen strains (66.7%) were susceptible (MIC ≤ 4 µg/mL), three (14.3%) were intermediate, and four (19.0%) were resistant (MIC ≥ 16 µg/mL). Non-susceptible strains, defined as intermediate or resistant, included M1, M3, M4, M5, M6, M8, and M22. An additional MBL-harboring strain, BAA-3302, was also resistant (MIC = 16 µg/mL).

**Table 1.** Antibiotic susceptibilities of each *A. baumannii* isolate to FDC. Strains are labeled M1-M22. Note that strain M15 was excluded from the study, as it was not *A. baumannii*. MIC values were determined at least two independent times, with the determinations listed as MIC1 and MIC2. Strains that are susceptible are denoted as S; strains with intermediate susceptibility are denoted as I; strains that are resistant are denoted as R, according to CLSI standards.

| Strain | MIC 1 | MIC 2 | S/I/R | Strain | MIC 1 | MIC 2 | S/I/R |
| --- | --- | --- | --- | --- | --- | --- | --- |
| M1 | 64 | 64 | R | M12 | 2 | 2 | S |
| M2 | 1 | 1 | S | M13 | 0.5 | 1 | S |
| M3 | 32 | 64 | R | M14 | 0.5 | 1 | S |
| M4 | 8 | 16 | I | M16 | 1 | 1 | S |
| M5 | 16 | 16 | R | M17 | 0.5 | 1 | S |
| M6 | 32 | 32 | R | M18 | 0.5 | 1 | S |
| M7 | 0.5 | 1 | S | M19 | 1 | 1 | S |
| M8 | 8 | 16 | I | M20 | 1 | 2 | S |
| M9 | 1 | 2 | S | M21 | 1 | 1 | S |
| M10 | 1 | 1 | S | M22 | 8 | 16 | I |
| M11 | 2 | 4 | S | BAA3302 | 16 | 16 | R |

### 2.2. Determination of the combinatorial effects of FDC with various antibiotics

To identify synergistic combinations, we screened 17 FDC-antibiotic pairings against the PDR strain M9 using the disc stacking method for screening. Eleven combinations showed increased inhibitions zones (>3 mm) and were selected for further evaluation via checkerboard assays (Table S1). We also tested FDC with SUL-DUR, eravacycline, and omadacycline, three recently introduced agents with activity against *A. baumannii* [22–24]. Lower MIC values observed in ID-CAMHB prompted adjustments in starting concentrations for SUL-DUR combinations (Table S2). These fourteen combinations were further evaluated for combinatorial effects using checkerboard assays. FICI values were calculated to determine whether an interaction was synergistic, additive, indifferent, or antagonistic. No combination was found to demonstrate antagonism. The lowest FICI values calculated for each combination against each strain are shown in Table 2, and overall rates of combinatorial effects for each antibiotic combination are shown in Table 3.

**Table 2.** Fractional inhibitory concentration index (FICI) values obtained from FDC in combination with various antibiotics against *A. baumannii* strains M1-M22. FICI values in the synergistic range (≤0.5) are reported in blue; FICI values in the additive range (0.5-1.0) are reported in yellow; FICI values indicating no interaction (1.0-4.0) are reported in pink.

| Paired Antibiotic | M1 | M2 | M3 | M4 | M5 | M6 | M7 | M8 | M9 | M10 | M11 | M12 | M13 | M14 | M16 | M17 | M18 | M19 | M20 | M21 | M22 |
| --- | --- | --- | --- | --- | --- | --- | --- | --- | --- | --- | --- | --- | --- | --- | --- | --- | --- | --- | --- | --- | --- |
| Ceftazidime | 1.01 | 0.56 | 0.51 | 0.19 | 0.28 | 1.50 | 0.13 | 0.56 | 1.01 | 0.16 | 0.14 | 1.01 | 1.01 | 0.25 | 1.06 | 0.75 | 0.63 | 0.28 | 0.52 | 1.06 | 1.01 |
| CAZ-AVI | 0.05 | 0.50 | 0.04 | 0.27 | 0.27 | 0.02 | 0.26 | 0.01 | 0.02 | 0.09 | 0.04 | 0.02 | 0.02 | 0.14 | 0.09 | 0.05 | 0.02 | 0.06 | 0.09 | 0.02 | 0.02 |
| Ceftriaxone | 0.63 | 0.51 | 0.51 | 0.53 | 0.26 | 1.01 | 0.27 | 0.14 | 0.27 | 0.51 | 0.51 | 1.01 | 0.14 | 1.01 | 0.26 | 1.00 | 0.75 | 0.51 | 0.27 | 0.14 | 0.19 |
| Cefotaxime | 0.53 | 0.53 | 1.01 | 0.51 | 0.63 | 0.25 | 0.16 | 0.16 | 0.27 | 0.53 | 0.56 | 1.002 | 1.02 | 0.25 | 1.00 | 0.63 | 1.06 | 0.56 | 1.03 | 1.06 | 0.75 |
| Cefepime | 0.31 | 0.28 | 0.38 | 0.38 | 0.56 | 0.50 | 0.56 | 0.75 | 0.51 | 0.53 | 0.19 | 0.51 | 0.09 | 0.75 | 0.53 | 0.75 | 0.31 | 0.50 | 0.51 | 0.63 | 0.31 |
| AMP-SUL | 0.14 | 1.06 | 0.25 | 1.25 | 1.06 | 0.25 | 1.06 | 1.02 | 0.31 | 0.56 | 0.75 | 0.28 | 0.04 | 0.56 | 1.12 | 1.12 | 1.12 | 0.75 | 0.75 | 0.75 | 0.28 |
| Sulbactam | 0.14 | 0.38 | 0.19 | 0.25 | 0.56 | 0.19 | 0.75 | 0.63 | 0.52 | 0.38 | 0.31 | 0.50 | 0.63 | 0.56 | 1.50 | 1.25 | 1.13 | 0.38 | 0.38 | 1.13 | 0.38 |
| SUL-DUR | 0.13 | 1.01 | 0.13 | 0.19 | 0.06 | 0.06 | 0.38 | 0.19 | 0.19 | 0.13 | 0.08 | 0.16 | 0.38 | 0.50 | 0.19 | 0.31 | 0.25 | 0.16 | 0.25 | 0.38 | 0.07 |
| Amikacin | 0.38 | 0.16 | 0.09 | 0.28 | 0.27 | 0.19 | 0.52 | 0.19 | 0.03 | 1.25 | 0.50 | 0.04 | 1.25 | 0.52 | 0.63 | 0.05 | 0.08 | 0.26 | 0.51 | 0.51 | 0.19 |
| Ciprofloxacin | 1.002 | 0.38 | 0.53 | 0.26 | 0.504 | 0.63 | 0.61 | 1.08 | 1.16 | 1.004 | 0.52 | 0.25 | 1.14 | 0.56 | 1.16 | 0.502 | 0.63 | 1.004 | 1.002 | 0.19 | 0.502 |
| Rifampin | 0.53 | 0.75 | 0.25 | 0.25 | 0.52 | 0.52 | 0.31 | 0.51 | 0.19 | 0.27 | 1.06 | 1.02 | 1.01 | 1.01 | 0.50 | 1.01 | 1.06 | 0.52 | 1.00 | 1.06 | 0.51 |
| Doxycycline | 0.504 | 0.13 | 1.002 | 2.00 | 2.00 | 0.13 | 0.25 | 0.31 | 0.25 | 0.13 | 0.14 | 0.38 | 0.19 | 0.13 | 0.53 | 0.75 | 3.00 | 0.51 | 0.06 | 0.13 | 0.28 |
| Omadacycline | 0.504 | 0.56 | 0.504 | 1.06 | 1.02 | 1.01 | 1.03 | 0.53 | 0.63 | 0.27 | 1.02 | 0.28 | 1.25 | 1.50 | 1.25 | 1.50 | 2.00 | 1.25 | 1.13 | 2.00 | 1.06 |
| Eravacycline | 0.75 | 1.06 | 0.50 | 0.75 | 0.50 | 0.53 | 1.00 | 1.06 | 0.50 | 0.63 | 1.00 | 0.63 | 1.02 | 0.75 | 1.06 | 1.50 | 1.06 | 0.63 | 0.75 | 1.06 | 1.00 |

**Table 3.** Rates of synergistic, additive, or lack of effects between antibiotics paired with FDC against strains M1-M22.

| Paired Antibiotic | Synergistic | Additive | Indifferent |
| --- | --- | --- | --- |
| Ceftazidime | 33.3% | 28.6% | 38.1% |
| CAZ-AVI | 100.0% | 0.0% | 0.0% |
| Ceftriaxone | 42.9% | 42.9% | 14.3% |
| Cefotaxime | 23.8% | 47.6% | 28.6% |
| Cefepime | 47.6% | 52.4% | 0.0% |
| AMP-SUL | 38.1% | 28.6% | 33.3% |
| Sulbactam | 52.4% | 28.6% | 19.0% |
| SUL-DUR | 95.2% | 0.0% | 4.8% |
| Amikacin | 66.7% | 23.8% | 9.5% |
| Ciprofloxacin | 19.0% | 42.9% | 38.1% |
| Rifampin | 23.8% | 42.9% | 33.3% |
| Doxycycline | 61.9% | 19.0% | 19.0% |
| Omadacycline | 9.5% | 19.0% | 71.4% |
| Eravacycline | 14.3% | 52.4% | 33.3% |

Interactions between antibiotic pairings were not uniform for all strains tested, in agreement with our previous findings.^21^ Several FDC-based combinations displayed synergy or additive effects across the 21-strain panel. Notably, FDC demonstrated the highest synergy rates in combination with CAZ-AVI (100.0%) and SUL-DUR (95.2%). FDC and amikacin showed synergy against 66.7% of strains, including all strains non-susceptible to FDC alone, and additive effects against 23.8% of strains. Doxycycline and FDC showed synergy against 61.9% of strains, and additive effects against 19.0% of strains. Sulbactam and FDC were also relatively effective, demonstrating synergy against 52.4% of strains, and additive effects against 28.6% of strains. FDC combined with other β-lactam drugs, including cefepime, ceftriaxone, cefotaxime, and AMP-SUL, showed moderate synergy rates ranging from 23.8-52.4%. Rifampin and ciprofloxacin displayed synergy in 19.0% and 23.8% of strains in combination with FDC, respectively, and additive effects against nearly half of strains. In contrast, FDC paired with omadacycline or eravacycline showed limited synergy (<15%), despite these drugs’ efficacy as monotherapy [21], although eravacycline and FDC showed additive effects against 52.4% of strains. No antagonism was observed with any combination. Representative checkerboard assays displaying synergy patterns are shown for six combinations (Figures 1-6), including FDC paired with CAZ-AVI, SUL-DUR, amikacin, doxycycline, AMP-SUL, and sulbactam. Representative checkerboard assays demonstrating synergy patterns for FDC combined with eravacycline, omadacycline, rifampin, ciprofloxacin, ceftazidime, cefepime, ceftriaxone, and cefotaxime are shown in Supplemental Figures S1-S8.

**Figure 1.**
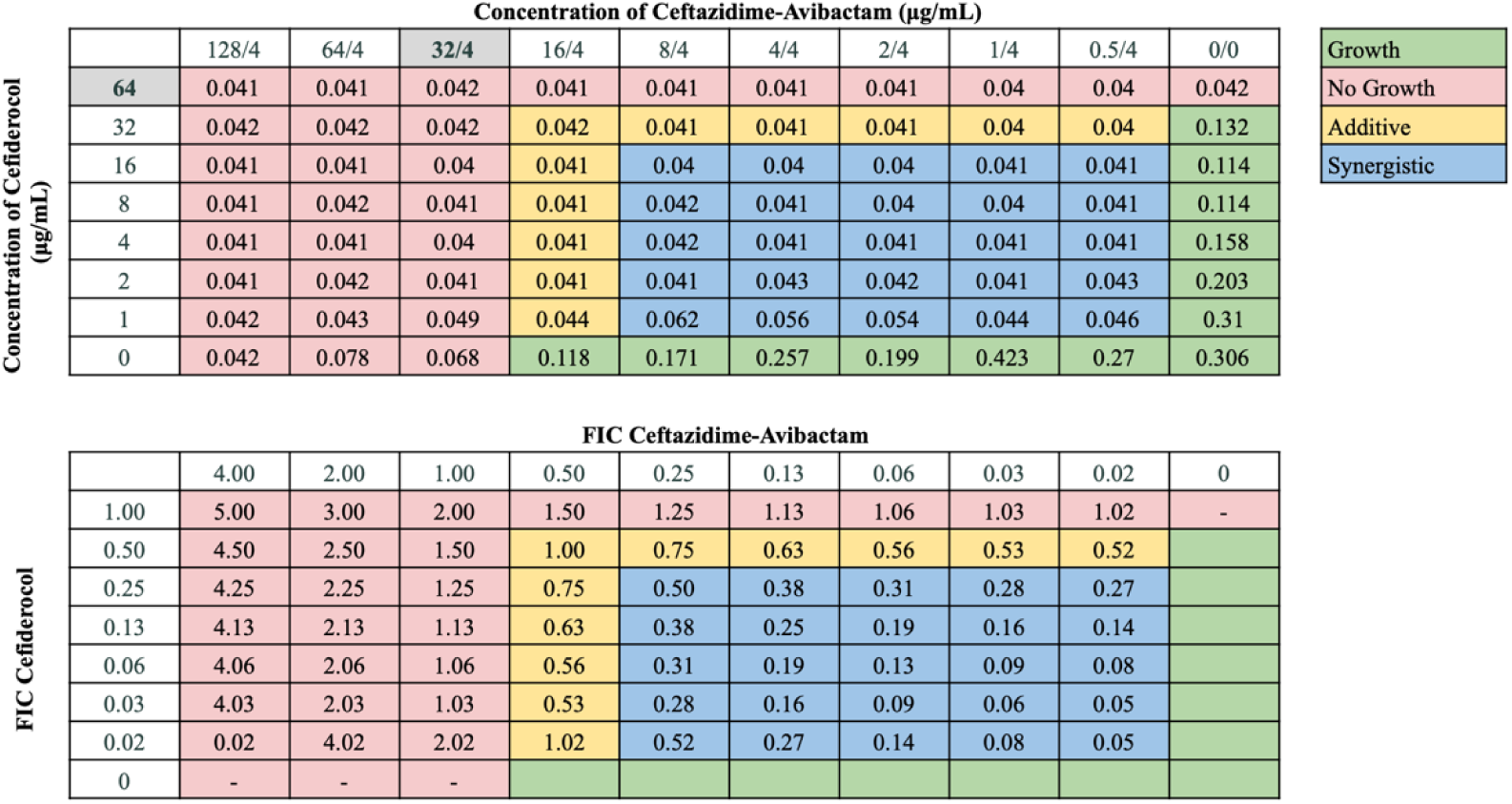
Representative checkerboard assay with FDC and CAZ-AVI against strain M1. Top: OD_600_ measurements following 16 hours of static growth at 37°C. The MIC values for each drug alone are highlighted in gray. The pink boxes indicate wells in which no bacterial growth occurred (OD_600_ <0.1), and green boxes indicate wells in which bacterial growth did occur. Bottom: FIC values were calculated for each drug (concentration/MIC) and added together for all wells where no growth was observed. The yellow boxes indicate additive interactions (FICI between 0.5-1.0), and blue boxes indicate synergistic interactions (FICI ≤0.5).

**Figure 2.**
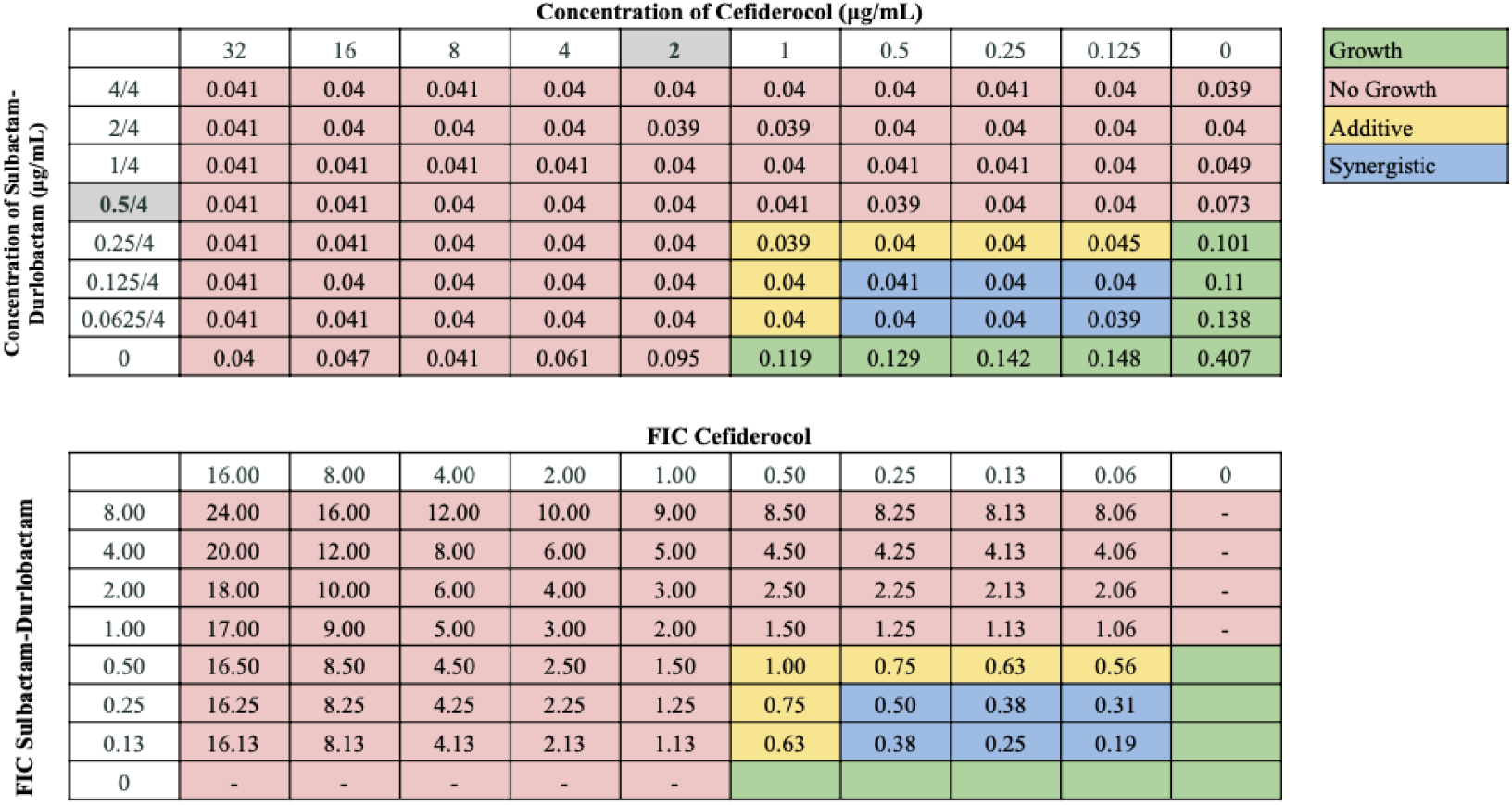
Representative checkerboard assay with FDC and SUL-DUR against strain M16. Top: OD_600_ measurements following 16 hours of static growth at 37°C. The MIC values for each drug alone are highlighted in gray. Note the low starting concentration and MIC values for SUL-DUR. The pink boxes indicate wells in which no bacterial growth occurred (OD_600_ <0.1), and green boxes indicate wells in which bacterial growth did occur. Bottom: FIC values were calculated for each drug (concentration/MIC) and added together for all wells where no growth was observed. The yellow boxes indicate additive interactions (FICI between 0.5-1.0), and blue boxes indicate synergistic interactions (FICI ≤0.5).

**Figure 3.**
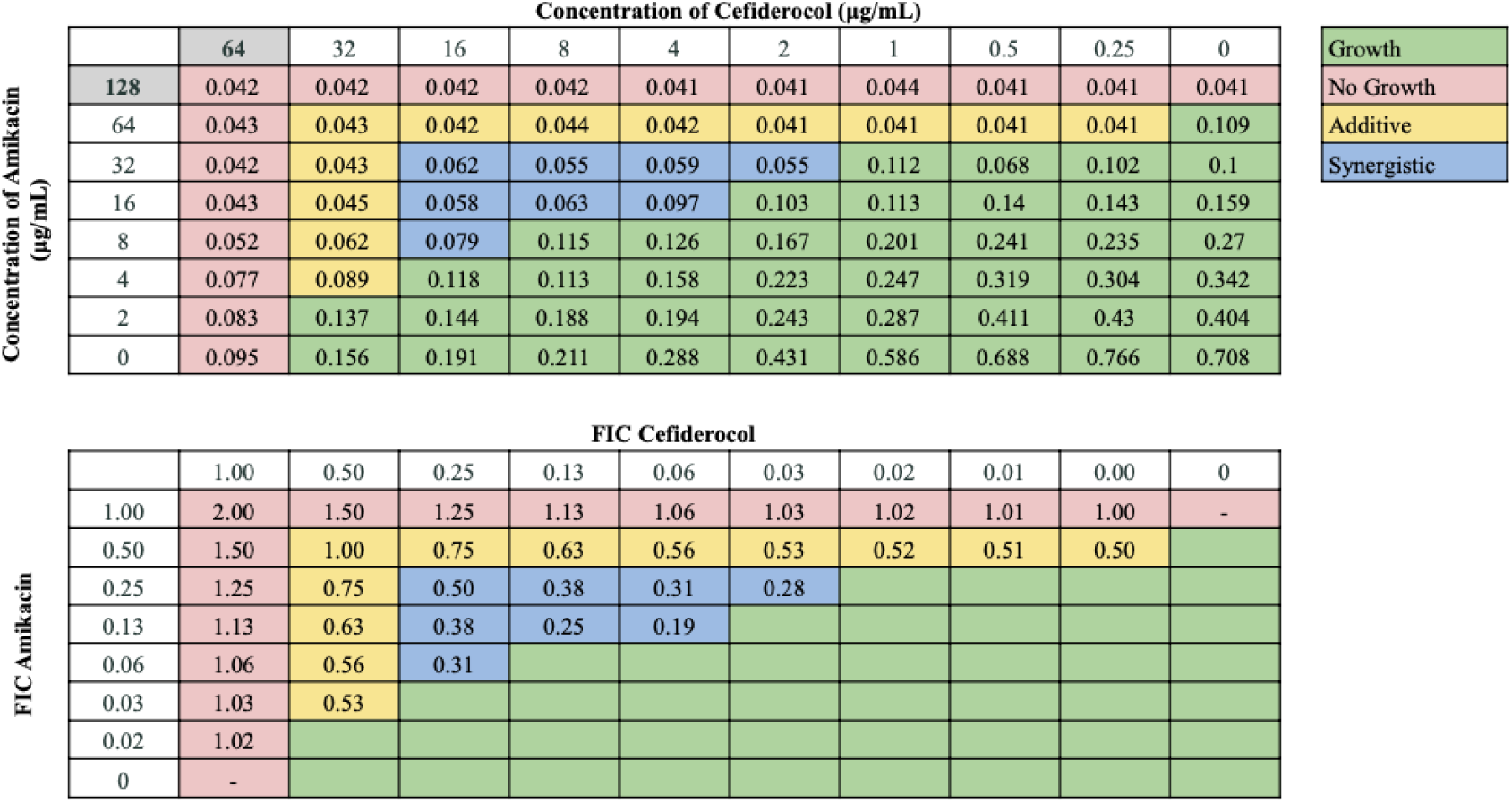
Representative checkerboard assay with FDC and amikacin against strain M6. Top: OD_600_ measurements following 16 hours of static growth at 37°C. The MIC values for each drug alone are highlighted in gray. The pink boxes indicate wells in which no bacterial growth occurred (OD_600_ <0.1), and green boxes indicate wells in which bacterial growth did occur. Bottom: FIC values were calculated for each drug (concentration/MIC) and added together for all wells where no growth was observed. The yellow boxes indicate additive interactions (FICI between 0.5-1.0), and blue boxes indicate synergistic interactions (FICI ≤0.5).

**Figure 4.**
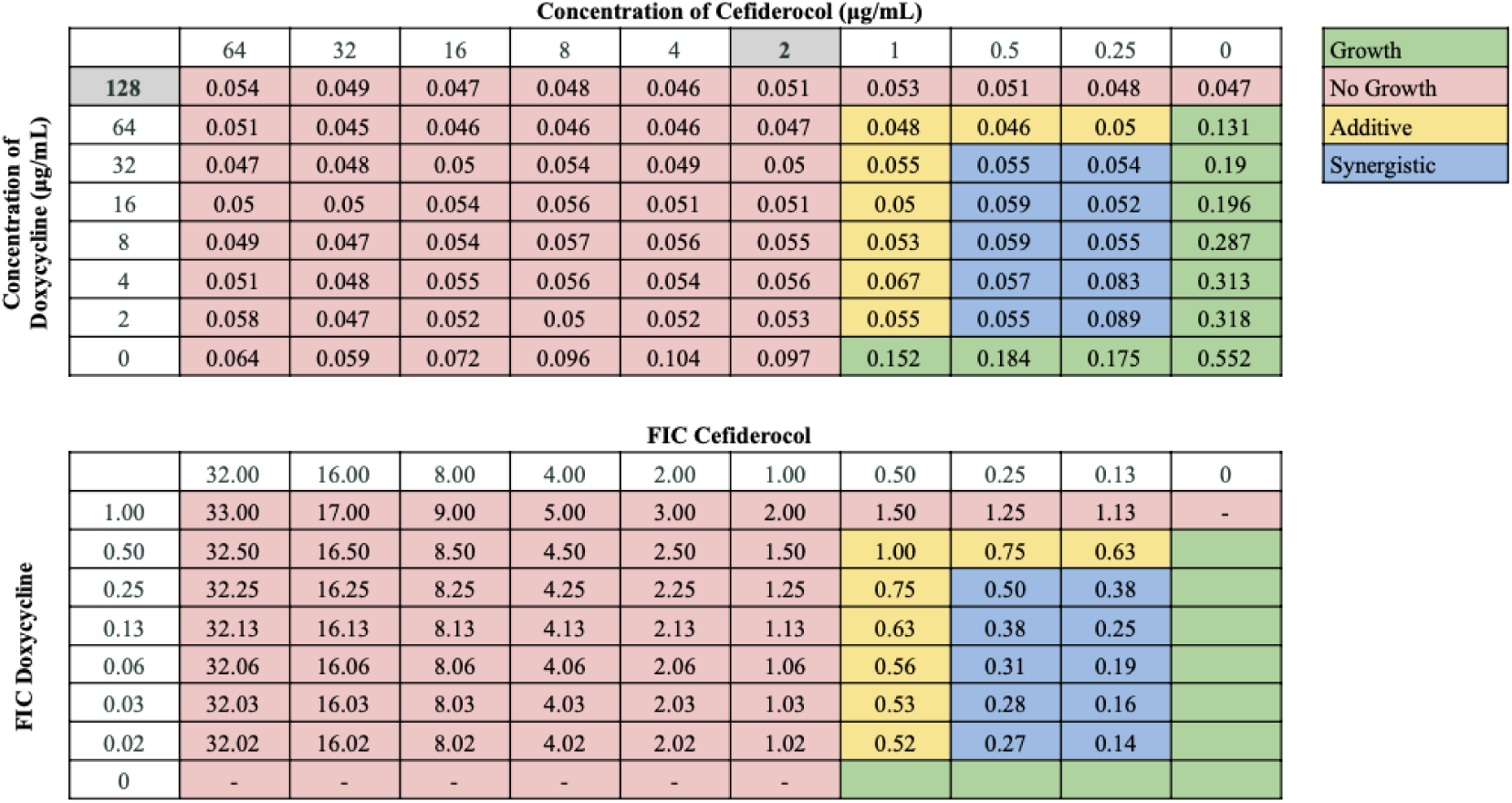
Representative checkerboard assay with FDC and doxycycline against strain M11. Top: OD_600_ measurements following 16 hours of static growth at 37°C. The MIC values for each drug alone are highlighted in gray. The pink boxes indicate wells in which no bacterial growth occurred (OD_600_ <0.1), and green boxes indicate wells in which bacterial growth did occur. Bottom: FIC values were calculated for each drug (concentration/MIC) and added together for all wells where no growth was observed. The yellow boxes indicate additive interactions (FICI between 0.5-1.0), and blue boxes indicate synergistic interactions (FICI ≤0.5).

**Figure 5.**
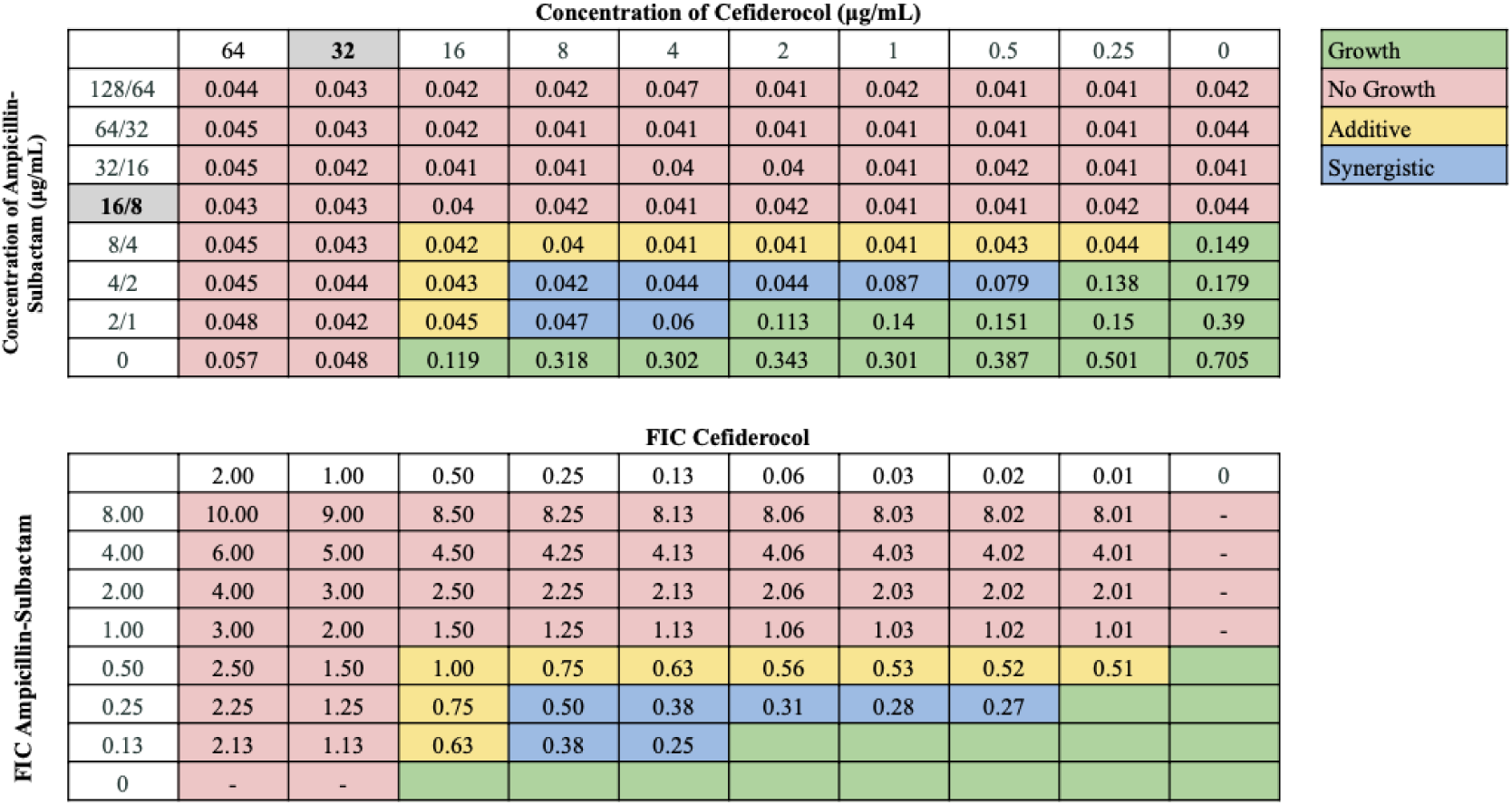
Representative checkerboard assay with FDC and AMP-SUL against strain M3. Top: OD_600_ measurements following 16 hours of static growth at 37°C. The MIC values for each drug alone are highlighted in gray. The pink boxes indicate wells in which no bacterial growth occurred (OD_600_ <0.1), and green boxes indicate wells in which bacterial growth did occur. Bottom: FIC values were calculated for each drug (concentration/MIC) and added together for all wells where no growth was observed. The yellow boxes indicate additive interactions (FICI between 0.5-1.0), and blue boxes indicate synergistic interactions (FICI ≤0.5).

**Figure 6.**
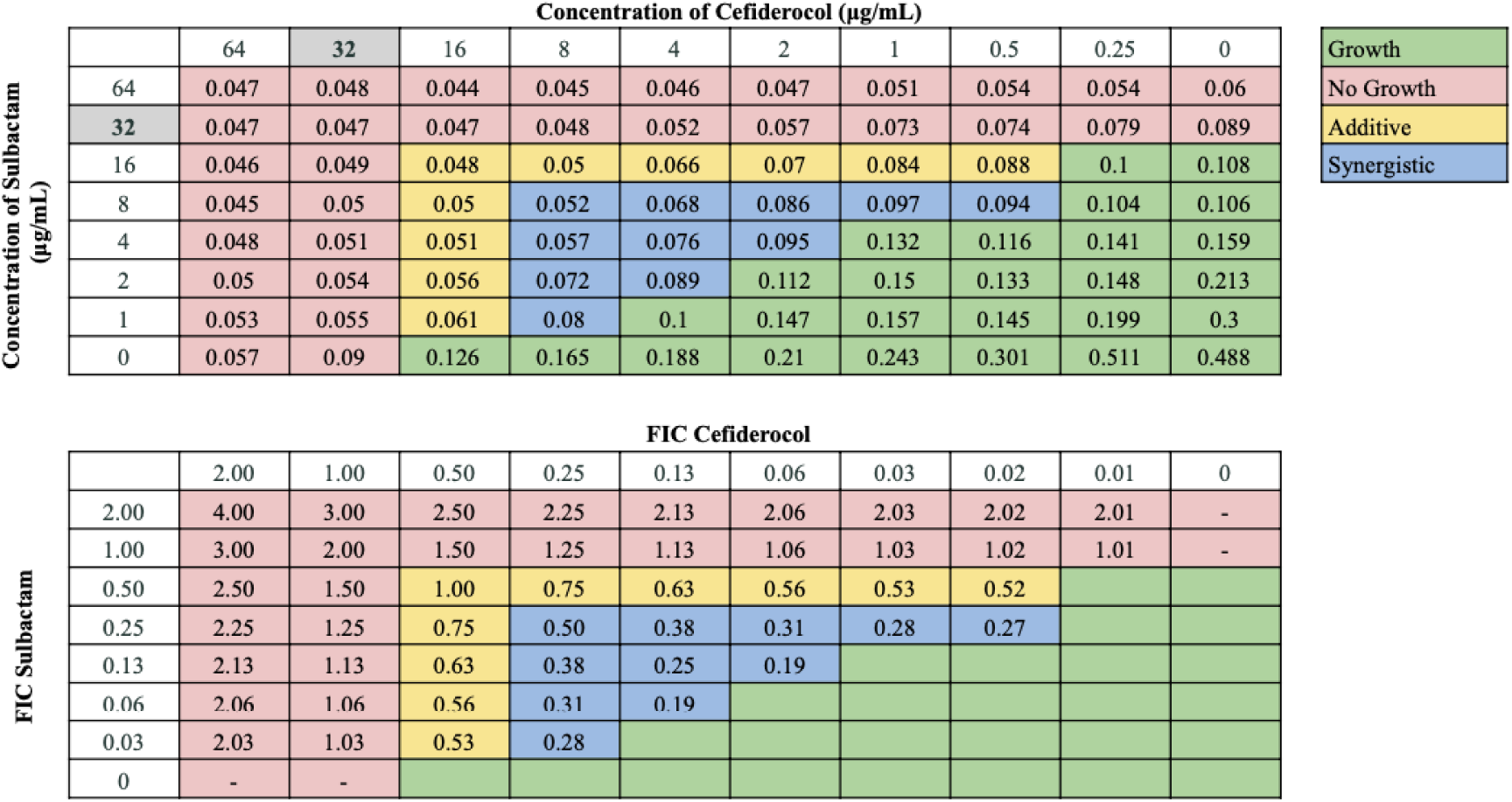
Representative checkerboard assay with FDC and sulbactam against strain M3. Top: OD_600_ measurements following 16 hours of static growth at 37°C. The MIC values for each drug alone are highlighted in gray. The pink boxes indicate wells in which no bacterial growth occurred (OD_600_ <0.1), and green boxes indicate wells in which bacterial growth did occur. Bottom: FIC values were calculated for each drug (concentration/MIC) and added together for all wells where no growth was observed. The yellow boxes indicate additive interactions (FICI between 0.5-1.0), and blue boxes indicate synergistic interactions (FICI ≤0.5).

### 3.3. Identification of resistance genes among FDC-resistant A. baumannii isolates

Previously, WGS was performed on eight selected strains, including three that were non-susceptible to FDC (M1, M4, M5) and five that were susceptible (M9, M10, M11, M13, M20) [21]. Resistance gene profiles are summarized in Table 4. All harbored class C (ADC type) β-lactamases, such as *blaADC-30, -33, -73, -80,* and *-150*. ADC-33 and ADC-73 appeared in both groups, while ADC-30, ADC-80, and ADC-150 were exclusive to susceptible strains. All strains also harbored class D (OXA) β-lactamases, including *blaOXA-23, -66, -83, -94,* and *-421*. OXA-82 was detected only in M1, an FDC-resistant strain. OXA-23 and OXA-66 were found across many sequenced isolates. OXA-94 was present in one non-susceptible and one susceptible strain. OXA-421 was limited to susceptible strains. Class B (MBL) β-lactamases were not found in our original collection of strains. A mutation in *ftsI* (PBP3) was also identified in both susceptible and non-susceptible isolates, suggesting that this is not a major determinant of FDC resistance. All strains carried aminoglycoside resistance determinants, including AME genes and *armA*, as well as *tetB* for tetracycline resistance and *gyrA* mutations linked to fluoroquinolone resistance.

**Table 4.**
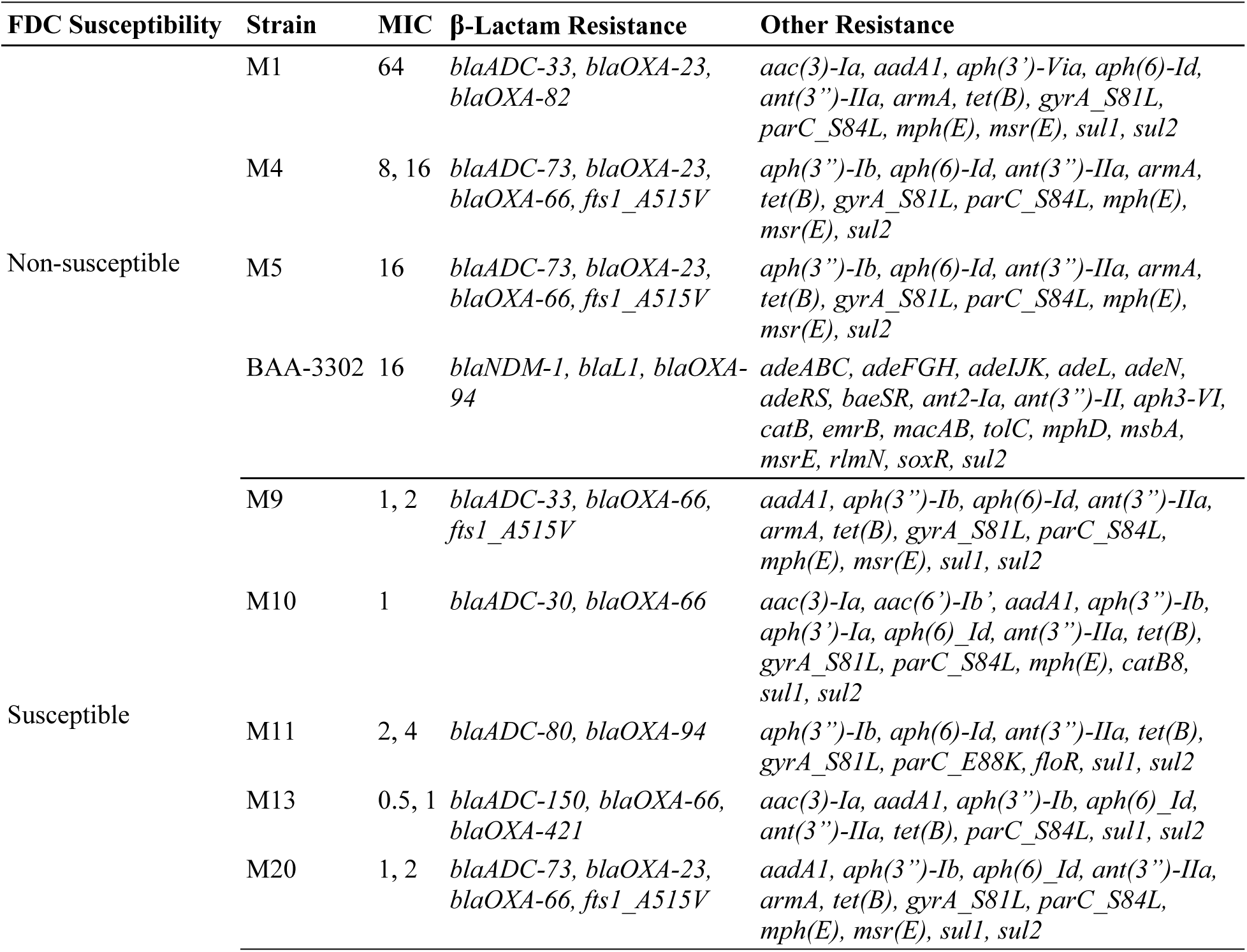
Identified resistance genes among nine selected *A. baumannii* strains [21]. Strains with FDC MIC values >4 are categorized as non-susceptible, while those with MIC values ≤4 are categorized as susceptible.

**Table 5.** Combinatorial effects of FDC-containing antibiotic combinations against strain BAA-3302, an *NDM-1*-harboring strain resistant to FDC (MIC = 16 μg/mL). FICI values are reported, as well as MIC values of FDC in the presence of each paired antibiotic.

| Paired Antibiotic | FICI Value | Combined FDC MIC ($\mu\text{g/mL}$) | Interpretation |
| --- | --- | --- | --- |
| Ceftazidime | 0.19 | 0.25 | Synergy |
| CAZ-AVI | 0.19 | 1 | Synergy |
| Ceftriaxone | 0.05 | 0.25 | Synergy |
| Cefotaxime | 0.03 | 0.25 | Synergy |
| Cefepime | 0.27 | 0.25 | Synergy |
| AMP-SUL | 0.14 | 0.25 | Synergy |
| Sulbactam | 1.01 | 16 | No Interaction |
| SUL-DUR | 0.19 | 1 | Synergy |
| Amikacin | 0.27 | 0.25 | Synergy |
| Ciprofloxacin | 0.03 | 0.25 | Synergy |
| Rifampin | 0.08 | 0.25 | Synergy |
| Doxycycline | 1.03 | 16 | No Interaction |
| Omadacycline | 1.03 | 16 | No Interaction |
| Eravacycline | 1.25 | 16 | No Interaction |

### 3.4. FDC-based combinations against an NDM-1-producing A. baumannii isolate

As our collection did not contain any MBLs, we tested the same antibiotic combinations against BAA-3302, an MBL-harboring clinical isolate that we determined to be resistant to FDC (Table 4). Notably, the β-lactam drugs ceftazidime, ceftriaxone, cefotaxime, and cefepime each reduced FDC’s MIC 64-fold, from 16 μg/mL to 0.25 μg/mL. FDC also synergized with all β-lactam/β-lactamase inhibitor combinations: AMP-SUL (MIC = 0.25 μg/mL), CAZ-AVI, and SUL-DUR (both MIC = 1 μg/mL). In addition, FDC showed strong synergy with amikacin, ciprofloxacin, and rifampin, each lowering its MIC to 0.25 μg/mL. No interaction was observed with doxycycline, omadacycline, or eravacycline. Representative checkerboard assay with FDC in combination with cefotaxime and ciprofloxacin are shown in Figures 7 and S9, respectively. The genome of strain BAA-3302 is freely available, and its resistance genes are cataloged (Table 4). Notably, BAA-3302 possesses two MBL genes, *blaNDM-1* and *blaL1*, as well as multiple RND-type efflux systems, such as *adeABC, adeFGH,* and *adeIJK*, which may affect susceptibility to a broad range of antibiotics, particularly tetracyclines.

**Figure 7.**
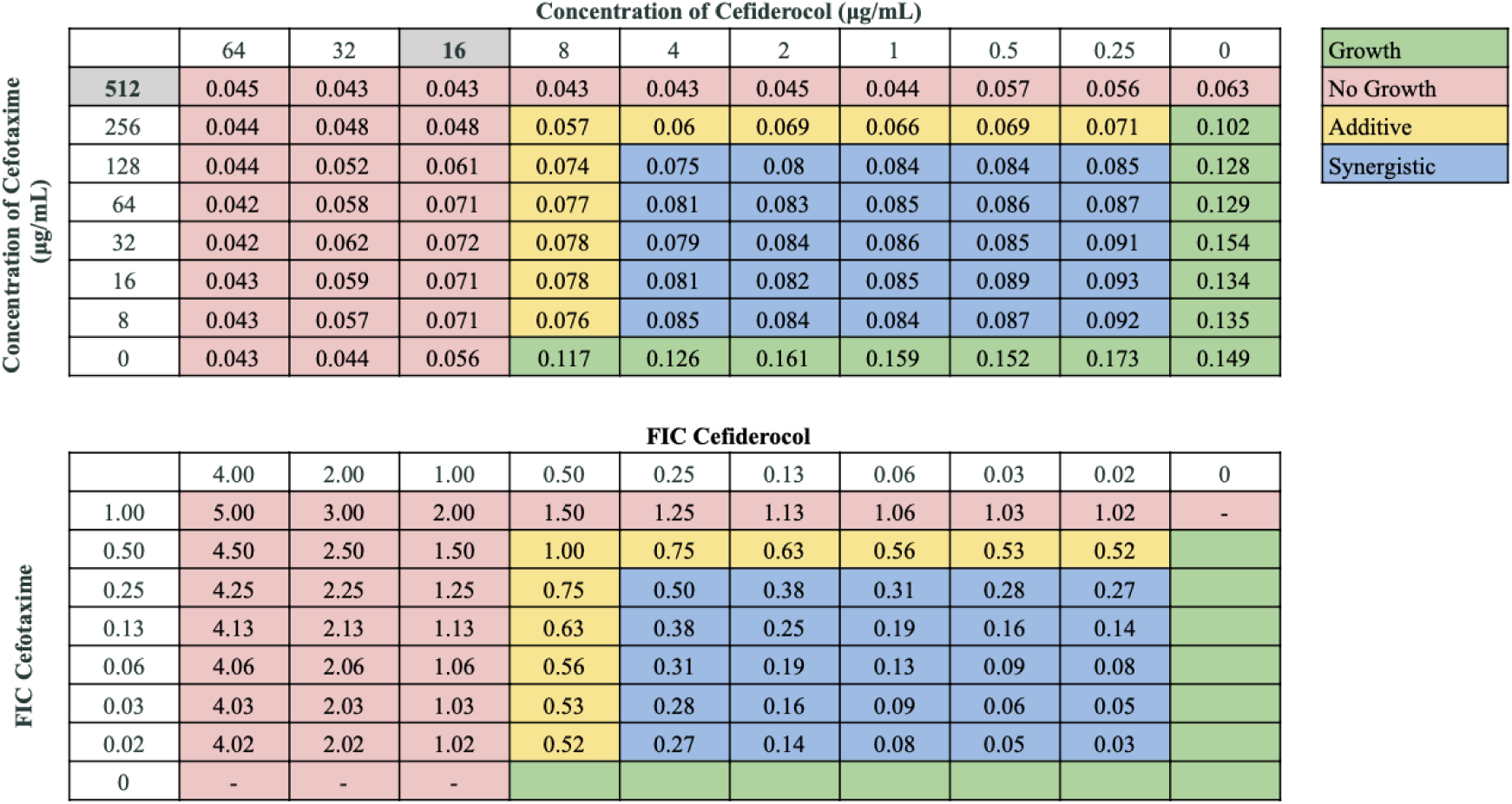
Representative checkerboard assay with FDC and cefotaxime against strain BAA-3302. Top: OD_600_ measurements following 16 hours of static growth at 37°C. The MIC values for each drug alone are highlighted in gray. The pink boxes indicate wells in which no bacterial growth occurred (OD_600_ <0.1), and green boxes indicate wells in which bacterial growth did occur. Bottom: FIC values were calculated for each drug (concentration/MIC) and added together for all wells where no growth was observed. The yellow boxes indicate additive interactions (FICI between 0.5-1.0), and blue boxes indicate synergistic interactions (FICI ≤0.5).

## 3. Discussion

### 3.1. FDC activity and resistance mechanisms

As antibiotic resistance rises in *A. baumannii*, treatment options become more limited. FDC has demonstrated both *in vitro* activity against CRAB and has been used clinically, although outcomes vary [13–15]. FDC features a catechol-type siderophore combined with a cephalosporin antibiotic containing side chains like those of ceftazidime and cefepime [11,25]. The catechol moiety chelates iron to facilitate uptake via iron transporter channels, a mechanism commonly referred to as the “Trojan Horse” mechanism [12]. FDC then dissociates from iron and inhibits PBP3 to disrupt peptidoglycan synthesis [12,25]. FDC’s unique mechanism and large structure confers increased stability against common antibiotic resistance mechanisms, particularly β-lactamases such as AmpC, ESBLs, OXA-type, and MBLs [25,26]. FDC demonstrated strong activity against our panel of 21 XDR and PDR *A. baumannii* strains, with 66.7% strains susceptible. Only four strains (M1, M3, M5, and M6), and the MBL-producing strain BAA-3302 were resistant. Known mechanisms of FDC resistance in *A. baumannii* include co-expression of multiple β-lactamases (e.g., NDM, KPC, OXA-type, and AmpC variants) and impaired siderophore-mediated uptake due to *tonB*-related loss of *piuA* and *pirA* expression [9,27].

WGS analysis revealed multiple β-lactamases across both susceptible and non-susceptible strains. M1, which had the highest FDC MIC value, was the only strain possessing *blaOXA-82*, suggesting a potential role for OXA-82 in FDC resistance. OXA-23, OXA-66, and OXA-94 were widely distributed across both susceptible and non-susceptible strains, suggesting they are insufficient alone to confer resistance but may modulate FDC activity when paired with other β-lactamases. Conversely, OXA-421 is unlikely to be a major resistance determinant, as it was only found in a susceptible strain. Class C cephalosporinases, such as ADC-33 and ADC-73, were detected in both susceptible and non-susceptible strains, but may contribute to resistance when co-expressed with OXA-type enzymes. Mutations in PBP3 (*ftsI*), the FDC target, were observed in both susceptible and non-susceptible strains, indicating they may not be a major mechanism of resistance. Strain BAA-3302 carried *blaNDM-1*, *blaL1*, and *blaOXA-94*. L1, a metallo-β-lactamase typically associated with *Stenotrophomonas maltophilia*, has broad activity against β-lactams and may contribute to FDC resistance similarly to NDM-1 [28]. Overall, FDC resistance appears multifactorial, involving multiple β-lactamases and target site alterations. However, as WGS was not performed on all isolates, additional genomic and functional studies are needed to comprehensively define resistance mechanisms.

### 3.2. Synergistic combinations including cefiderocol

Clinical trial outcomes of FDC for treating *A. baumannii* infections have been mixed. The CREDIBLE-CR trial reported that patients with CRAB infections treated with FDC demonstrated similar rates of infection resolution, but higher mortality rates compared to patients treated with the best available therapy, which usually included colistin-based regimens [16]. However, this higher mortality rate may have been due to baseline differences in illness severity [16]. Conversely, the APEKS-NP trial found that FDC was non-inferior to meropenem for nosocomial pneumonia, although *A. baumannii* infections only comprised 16% of patients studied [17]. With strong clinical data lacking, the IDSA recommends FDC only when first-line options such as SUL-DUR are unavailable, and always in combination [18]. However, few FDC-containing antibiotic combinations effective against *A. baumannii* have been reported. In a study of two CRAB isolates, Palombo et al. demonstrated that FDC in combination with fosfomycin yielded synergy against one strain and found no interaction between FDC and either piperacillin-tazobactam, CAZ-AVI, imipenem-relebactam, meropenem-vaborbactam, or AMP-SUL [29]. Another recent study reported *in vitro* synergy between FDC and either avibactam, sulbactam, meropenem, or amikacin against *A. baumannii* isolates [30]. Overall, we report multiple antibiotics that demonstrate potent synergy when combined with FDC.

Among the combinations tested, CAZ-AVI and SUL-DUR demonstrated the most consistent and potent synergy with FDC, with synergy observed in 100% and 95.2% of strains, respectively. These combinations significantly lowered FDC MICs, including in strains with high baseline resistance. The high synergy rate with both drugs are likely due to the broad spectrum inhibition of classes A, C, and D β-lactamases by avibactam and durlobactam, to protect the β-lactam components from degradation [31,32]. Moreover, additional targeting of PBP3 by ceftazidime, sulbactam, and durlobactam may further complement FDC activity [33,34]. FDC in combination with either CAZ-AVI or SUL-DUR present highly promising combination therapy options for drug-resistant *A. baumannii* infections. The strong synergy demonstrated between FDC and SUL-DUR is clinically relevant, as SUL-DUR is a new β-lactam/β-lactamase inhibitor combination now recommended by the IDSA as first-line therapy for CRAB, after the ATTACK trial demonstrated SUL-DUR’s non-inferiority to colistin in treating CRAB infections [18,22]. SUL-DUR was highly effective against our collection as a monotherapy, with 100.0% of strains susceptible, as defined as MIC values ≤4/4 [35]. The combination of FDC and SUL-DUR may be a highly promising alternative in combating highly drug-resistant *A. baumannii*, and further clinical evaluation should be conducted.

Among the standard-of-care drugs, amikacin showed the highest synergy rate (66.7%) with FDC, and lowered FDC MICs up to 32-fold in all strains non-susceptible to FDC. This synergy occurred despite widespread aminoglycoside resistance genes (AMEs, *armA*), likely due to complementary mechanisms [21,36]. It should be noted that aminoglycoside-β-lactam combination therapy is associated with higher mortality and adverse effect rates than β-lactam monotherapy, so while amikacin and FDC may an effective combination, their clinical use should be carefully weighed [37]. Doxycycline and FDC also showed high synergy rates (61.9%). Our collection was universally resistant to doxycycline, likely due to widespread distribution of the TetB efflux pump [21]. However, FDC paired with either omadacycline or eravacycline, two new tetracycline derivatives, did not exhibit high synergy rates, although eravacycline showed moderate additive effects with FDC. Omadacycline and eravacycline are effective against our collection as monotherapies [21]. Furthermore, no combinatorial effects with FDC in combination with either minocycline or tigecycline during initial disk stacking screening were observed, although minocycline does possess strong *in vitro* activity against our collection [21]. Our findings suggest doxycycline may be effective in combination with FDC against *A. baumannii*. Other non-β-lactam antibiotics showed variable efficacy. Rifampin in combination with FDC was relatively effective, with synergy against 5 strains (23.8%) and additive effects against 9 strains (42.9%). While rifampin has shown *in vitro* synergy with colistin [38], clinical trials have not demonstrated improved outcomes [39,40], and the IDSA does not recommend its use for CRAB [18]. However, our findings suggest rifampin with FDC may be effective against certain highly drug-resistant strains. Ciprofloxacin also demonstrated modest synergy (19.0%) with FDC, and additive effects against 42.9% of strains, despite universal resistance conferred by *gyrA* and *parC* mutations [21,41]. Interactions between FDC and non-β-lactam drugs may be due to improved drug uptake in the setting of FDC-mediated disruption of cell wall integrity [42].

Other cephalosporins demonstrated variable synergy with FDC. Synergy was observed with cefepime (47.6%), ceftriaxone (42.9%), ceftazidime (33.3%), and cefotaxime (23.8%). The differences in synergy among different cephalosporins may reflect differences in PBP3 targeting, although further research characterizing their exact mechanisms are needed. Sulbactam had higher synergy rates than AMP-SUL when combined with FDC (52.4% vs. 38.1%), suggesting the β-lactam component may not meaningfully enhance synergy with FDC. Since ampicillin has a broad range of PBP targets in *Acinetobacter*,^43^ while both FDC and sulbactam target PBP3 [11,25,32] we expected increased synergy between FDC and AMP-SUL due to increased PBP targets and additional β-lactamase protection from sulbactam. However, our findings suggest that FDC and sulbactam alone was a more effective combination, warranting further investigation into the underlying mechanisms of their synergy. Notably, no antagonism was observed in any combination, suggesting that FDC can be paired with a wide range of antibiotics. Overall, the most promising combinations, CAZ-AVI, SUL-DUR, amikacin, and doxycycline, demonstrated consistent synergy across numerous strains and are strong candidates for *in vivo* or clinical evaluation.

### 3.3. Cefiderocol synergism against an MBL-producing strain

While SUL-DUR is highly effective against *A. baumannii* and now recommended as first-line therapy, resistance has emerged, mainly due to NDM-type MBLs [44,45]. MBL-producing *A. baumannii* pose a major therapeutic challenge, as they also confer resistance to many other β-lactam drugs including carbapenems and FDC [46,47]. MBLs are not inhibited by durlobactam and avibactam due to their zinc-based active site [31,32,46]. None of the strains in our original collection (M1-M22) harbor MBLs, and all were susceptible to SUL-DUR [21]. To evaluate activity against MBL-producing strains, we tested antibiotic combinations against BAA-3302, an MBL-carrying strain resistant to both FDC and SUL-DUR. FDC combined with β-lactams or beta-lactamase inhibitor combinations, including ceftriaxone, cefotaxime, cefepime, ampicillin/sulbactam, CAZ-AVI, and SUL-DUR, showed strong synergy. Although MBLs confer some resistance to FDC, the drug is still relatively stable to MBL hydrolysis and has a weak binding affinity for the MBL active sites. This may allow FDC to enhance the activity of other β-lactams to overcome MBL-mediated degradation [25,47]. Additionally, the non-β-lactam drugs amikacin, ciprofloxacin, and rifampin displayed strong synergy with FDC against strain BAA-3302. This synergy may also be due to FDC-mediated inhibition of peptidoglycan synthesis which may compromise cell wall integrity, facilitating increased uptake of these agents [42]. These findings suggest potential combination strategies for treating highly resistant, MBL-producing *A. baumannii*.

No synergy was observed between FDC and tetracyclines, including eravacycline, omadacycline, and doxycycline, against BAA-3302. BAA-3302 was highly susceptible to tetracyclines, likely due to the absence of typical tetracycline resistance genes such as *tetB*. However, BAA-3302 possesses multiple RND efflux pumps not found in M1-M22, including *adeABC, adeFGH,* and *adeIJK*, which extrude a wide range of antibiotics and have particularly high activity against tetracyclines [48]. Active efflux of tetracyclines by RND pumps may have offset any increased membrane permeability conferred by FDC, thus limiting intracellular tetracycline accumulation and preventing synergistic activity [48]. In contrast to BAA-3302, many strains in our M1–M22 collection exhibited synergy between FDC and tetracyclines and other non-β-lactam antibiotics, including amikacin, ciprofloxacin, and rifampin, possibly due to the absence of *adeABC, adeFGH*, and *adeIJK*. A key limitation of this study is the inclusion of only a single MBL-harboring strain. Future studies should evaluate a broader panel of MBL-producing isolates with diverse resistance gene profiles to better characterize FDC-based combination efficacy. Overall, our findings may help guide decision-making in selecting additional antibiotics to combine with FDC in the treatment of highly drug-resistant *A. baumannii* infections.

## 4. Materials and Methods

### 4.1. Bacterial Strains, Media, and Growth Conditions

Strains M1-M22 are 21 *A. baumannii* isolates were described and characterized previously [21]. Strain BAA-3302 is a drug-resistant strain that harbors multiple MBLs, including NDM-1, purchased from ATCC. Iron-depleted, cation-adjusted Mueller Hinton broth (ID-CAMHB) was prepared as described previously [49]. Bacterial strains were inoculated into ID-CAMHB and grown overnight in a 37°C incubator with shaking. Bacterial growth was then assessed by measuring the optical density at 600 nm (OD_600_). FDC was provided by Shionogi (Florham Park, NJ, USA). Omadacycline was provided by Paratek Pharmaceuticals (Boston, MA, USA). Durlobactam was provided by Innoviva (Burlingame, CA, USA). All other antibiotics were purchased as previously described [50].

### 4.2. Determination of the MIC by Broth Microdilution

The MIC values of FDC against each strain were determined using broth microdilution according to standard protocols freely available from the Clinical & Laboratory Standards Institute (CLSI). Following overnight growth, cells were diluted into ID-CAMHB at a starting OD_600_ value of 0.05. FDC was added to each first well of a flat-bottom 96-well plate at a 2X concentration, with an “X” starting value of 64 μg/mL. Two-fold serial dilutions were performed along each row, and an equal volume of diluted bacterial cells were added to each well. The plates were incubated overnight in a 37°C incubator without shaking. The OD_600_ values were read using a Synergy H1 Microplate reader (Biotek) the next day. MIC determinations were made for at least two independent biological replicates. CLSI breakpoint data were used to determine the antimicrobial susceptibility status of the *A. baumannii* strains against FDC. The CLSI considers a MIC of ≤4 to be susceptible, >4 and <16 to be intermediate, and ≥16 to be resistant [50]. We noted from the checkerboard assays with SUL-DUR and FDC that the MIC values were 4- to 16-fold lower in ID-CAMHB. Therefore, MIC values were determined for SUL-DUR in both Cation-adjusted Mueller-Hinton Broth (CAMHB) and ID-CAMHB to guide appropriate “X” starting values for SUL-DUR-FDC checkerboard assays.

### 4.3. Screening for Combinatorial Effects by Disc Stacking Assays

Disc stacking assays were performed to screen for potential synergistic or additive interactions between FDC and other antibiotics, as previously described [51]. Overnight cultures were diluted into 2 mL of fresh ID-CAMHB and allowed to grow at 37°C for 2 to 3 hours. We selected strain M9, a PDR strain, as a representative strain for these disc stacking assays. Following this pre-growth phase, 300 μL of culture was spread onto a 150 mm x 15 mm MHA plate. Commercially available discs containing tetracycline, trimethoprim-sulfamethoxazole, levofloxacin, ceftriaxone, and piperacillin-tazobactam were used. For all other antibiotics tested (FDC, meropenem, ampicillin-sulbactam (AMP-SUL), sulbactam, amikacin, minocycline, tigecycline, and rifampin), filter paper discs 6 mm in diameter were prepared by adding a specific amount of drug according to CLSI recommendations, as previously described, and allowing them to dry for at least one hour at 30°C [49]. Plates were prepared evaluating FDC in combination with each of the antibiotics. Four antibiotic discs, two impregnated with FDC and two impregnated with a different antibiotic, were placed on the surface of the agar, evenly spaced from each other. A disc impregnated with FDC was then placed on top of one of the discs containing the other antibiotic, and a disc impregnated with the other antibiotic was placed on top of one of the FDC discs. 30 μL of sterile phosphate buffered saline was added to the top of the stacked discs. Plates were then incubated overnight in a 37°C incubator, and the zones of inhibition were measured the following day. A larger zone of inhibition of >3 mm surrounding the stacked discs compared to each drug alone indicated a potential synergistic or additive interaction, which was then further evaluated by the checkerboard assay.

### 4.4. Checkerboard Assays

Checkerboard assays were performed as previously described [49]. Two-fold serial dilutions of FDC and a different antibiotic were performed horizontally and vertically in a 96-well plate. Antibiotics were initially added at a 4X concentration, using the same “X” starting values as previously described [50]. For checkerboard assays involving SUL-DUR, sulbactam was added at an initial 4X concentration of 32 μg/mL and serial dilutions were then performed, while durlobactam was added at a constant concentration of 4 μg/mL per well, per CLSI recommendations [35]. Similarly for checkerboard assays involving ceftazidime-avibactam (CAZ-AVI), ceftazidime was added at an initial 4X concentration with subsequent serial dilutions performed, while avibactam was added at a constant concentration of 4 μg/mL per well [35]. Diluted cells were then added at an OD_600_ of 0.05 and incubated overnight. The OD_600_ was read the following morning. Combinatorial effects were measured by calculating the fractional inhibitory concentration index (FICI). The fractional inhibitory concentration (FIC) for each drug was determined, which is the concentration of the antibiotic in combination divided by the MIC of the antibiotic alone. The FICI was then calculated using the following formula: FICI = (MIC A_A+B_/MIC A) + (MIC B_A+B_/MIC B), in which MIC A and MIC B denote the MIC value of each antibiotic alone and MIC A_A+B_ and MIC B_A+B_ denote the MIC values of the drugs in combination. For a FICI of ≤0.5, the combination is synergistic; for a FICI of 0.5-1.0, the combination is additive; for a FICI >1 and <4, there is no effect; for a FICI ≥4, the interaction is antagonistic. All determinations were made at least two independent times.

## Acknowledgements

We thank Paratek Pharmaceuticals (Boston, MA) for providing omadacycline powder, Shionogi (Florham Park, NJ, USA) for providing Cefiderocol powder, and Innoviva (Burlingame, CA, USA) for providing durlobactam and eravacycline powders.

## Notes

### Competing Interest Statement

The authors have declared no competing interest.

